# Sex-specific Effects of SGLT2 Inhibition in a Kidney with Reduced Nephron Number: Modeling and Analysis

**DOI:** 10.1101/2021.12.16.472644

**Authors:** Anita T. Layton

## Abstract

The kidney plays an essential role in regulating the homeostasis of electrolytes, acid-base species, and fluids. Kidney structure and function are significantly affected in diabetes. These pathophysiological changes include glomerular hyperfiltration and tubular hypertrophy, and ultimately leading to diabetic kidney disease. A class of medications that have shown promise in slowing the progression to diabetic kidney disease are the sodium-glucose cotransporter 2 (SGLT2) inhibitors. SGLT2 inhibitors target Na^+^ and glucose reabsorption along the proximal convoluted tubule, enhance urinary glucose, Na^+^ and fluid excretion, and lower hyperglycemia in diabetes. We postulate that both diabetes-induced and SGLT2 inhibition-induced changes in kidney may exhibit significant sex differences, because the distribution of renal transporters along the nephron may be markedly different between women and men, as recently shown in rodents. The goals of this study are to (i) analyze how kidney function is altered in male and female patients with diabetes, (ii) assess the renal effects, in women and men, of an anti-hyperglycemic therapy that inhibits the sodium-glucose cotransporter 2 (SGLT2) in the proximal convoluted tubules, and (iii) study how those renal effects are altered in uninephrectomy. To accomplish these goals, we have developed computational models of kidney function, separate for male and female patients with diabetes and/or uninephredctomy. The simulation results indicate that by inducing osmotic diuresis in the proximal tubules, SGLT2 inhibition reduces paracellular transport, eventually leading to diuresis and natriuresis.

## Introduction

Diabetes affects >450 million people worldwide.^1,2^ Elevated blood glucose caused by diabetes can lead to diabetic kidney disease (DKD), which increases the risk of end-stage kidney disease (ESKD) requiring dialysis or a kidney transplant. DKD also promotes cardiovascular disease, the leading cause of mortality in persons with DKD. A crucial ingredient in developing effective therapies to halt or slow the progression of diabetes and DKD is to understand how the disease and therapies may affect men and women differently. Indeed, sex differences have been reported in cardiovascular disease, autoimmune disease, respiratory disease, and cognitive disease; diabetes and DKD are no different. Overall, women appear to be less prone to developing diabetes, although diabetic comorbidities including cardiovascular disease and ESKD are more likely to occur in women than in men.

The sex difference in the development of DKD and cardiovascular disease in patients with diabetes is not unexpected. Sex hormones are known to regulate not only the reproductive system, but the structure and function of nearly every tissue and organ in the mammalian body. In particular, the kidney exhibits notable sex differences in its function. ^3,4^ The functional implications of the sexual dimorphism in kidney morphology, hemodynamics, and transporter pattern have been investigated in recent studies.^5-8^ Given the kidney’s key role in regulating electrolyte and volume balance, sex differences in blood pressure control and hypertension are well known.^9,10^ The kidney undergoes major changes as diabetes progresses, including hypertrophy, hyperfiltration, and alterations in transporter expression. Given the sex differences in renal structure and function, to what extent are diabetes-induced alterations in kidney function different between the sexes?

For decades, DKD treatment has focused on glycemic and blood pressure control, the latter especially with renin-angiotensin-aldosterone system blockers.^1,2^ More recently, medications called sodium-glucose cotransporter-2 (SGLT2) inhibitors have become part of standard of the care for improving glycemic control by promoting urine glucose excretion. SGLT2 inhibitors differs from other popular anti-hyperglycemic medications in that they target no the insulin pathway but the kidneys. Besides the excretion of metabolic wastes and the maintenance of electrolyte and acid-base balance, the kidneys also play a key role in regulating blood pressure and blood glucose. As such, SGLT2 inhibitors reduce attenuate postprandial increases in blood glucose by targeting its reabsorption along the early proximal tubule and enhancing urinary glucose excretion. Because the transport of glucose and Na^+^ in the proximal tubule is coupled, inhibition of SGLT2 also reduces proximal tubular Na^+^ and fluid reabsorption, and induces natriuresis and diuresis. Hence, in addition to its anti-hyperglycaemic effect, SGLT2 inhibitors have been reported to reduce heart failure and cardiovascular events by 20-40% in patients with T2D.^11,12^

While SGLT2 inhibitors have shown much promise, many aspects, regarding its potential side-effects and mechanisms, remain unclear. One important but insufficiently understood is its sex-specific impact on kidney function. Under healthy conditions, higher expressions of SGLT1 and SGLT2 have been observed in female rats compared with male rats.^3,13^ Inhibiting SGLT2 lowers the proximal tubule uptake of not only glucose but Na^+^ as well, thereby shifting glucose and Na^+^ reabsorption to downstream nephron segments. This effect deserves careful examination for two reasons. First, the efficiency of Na^+^ transport (namely, the number of Na^+^ moles reabsorbed per O_2_ moles consumed) changes along the nephron, with the early proximal tubule being more efficient than the distal segments.^14^ To what extent renal tubular oxygen consumption change due to SGLT2 inhibition, and how does that change differs between men and women, and between a healthy and diabetic kidney?

Furthermore, sex dimorphism has been reported not only in SGLT2 expression level but other renal transporters as well. Findings by Veiras et al. ^15^ indicate that, compared with male rat nephrons, female rat nephrons exhibit, in the proximal tubule, greater Na^+^/H^+^ exchanger 3 (NHE3) phosphorylation a higher distribution of NHE3 at the base of the microvilli, and lower abundance of Na^+^-P_i_ cotransporter 2 (NaPi2), aquaporin-1 (AQP1), and claudin-2. These differences are associated with lower Na^+^ and HCO3 ^-^ reabsorption and increased volume flow from the proximal tubule.^15^ In contrast, transporters along the distal nephrons exhibit higher abundance of total and phosphorylated Na^+^-Cl^-^ cotransporter (NCC), claudin-7, and cleaved forms of epithelial Na^+^ channel (ENaC) α-and γ-subunits in female rats.^15^ How might the sex differences in renal transporter pattern affect the kidney’s response to SGLT2 inhibition?

Finally, SGLT2 inhibitors have also been approved in patients with DKD. When a substantial fraction of the nephron population is lost, the remaining functional nephrons in these patients undergo adaptations including hyperfiltration and increase in transport capacity. Thus, these kidneys may be particularly sensitive to changes in transport work caused by SGLT2 inhibition and to developing hypoxia. To better understand the nuances and long-term effects of SGLT2 inhibitors, we sought to assess the extent to which diabetes and SGLT2 inhibition can affect transport activity and renal metabolism in a nephrectomized kidney.

## Modeling Methodology

Model simulations were performed using our published model of epithelial transport along human nephrons.^5-8,16-25^ The model represents flow-dependent transport along the proximal tubule, connecting tubule, and cortical collecting duct. Sex-specific model parameters are used, which are then further adjusted from baseline to simulate a kidney in (i) male and female patients with diabetes, (ii) male and female patients with uninephrectomy (UNX), and (iii) male and female patients with diabetes and UNX.

## Simulation Results

### Kidney function in non-diabetic women and men under SGLT2 inhibition

The model simulates SGLT2 inhibition by blocking 90% of its activity. In a non-diabetic kidney, the remaining 10% of the SGLT2 mediates the reabsorption of ∼30% of the filtered glucose along the proximal convoluted tubules. The S3 segment’s SGLT1-mediated glucose transport is consequently increased to ∼40% of filtered load. The osmotic diuretic effect of the SGLT2 inhibition reduces passive paracellular reabsorption in the proximal tubule. Solute transport shifts downstream, primarily to the medullary thick ascending limbs, but also to the distal segments (Figs. 1-3). The increase in thick ascending limb Na^+^ transport is larger in women compared to men, resulting in more severe natriuresis in men.

**Figure 1.**
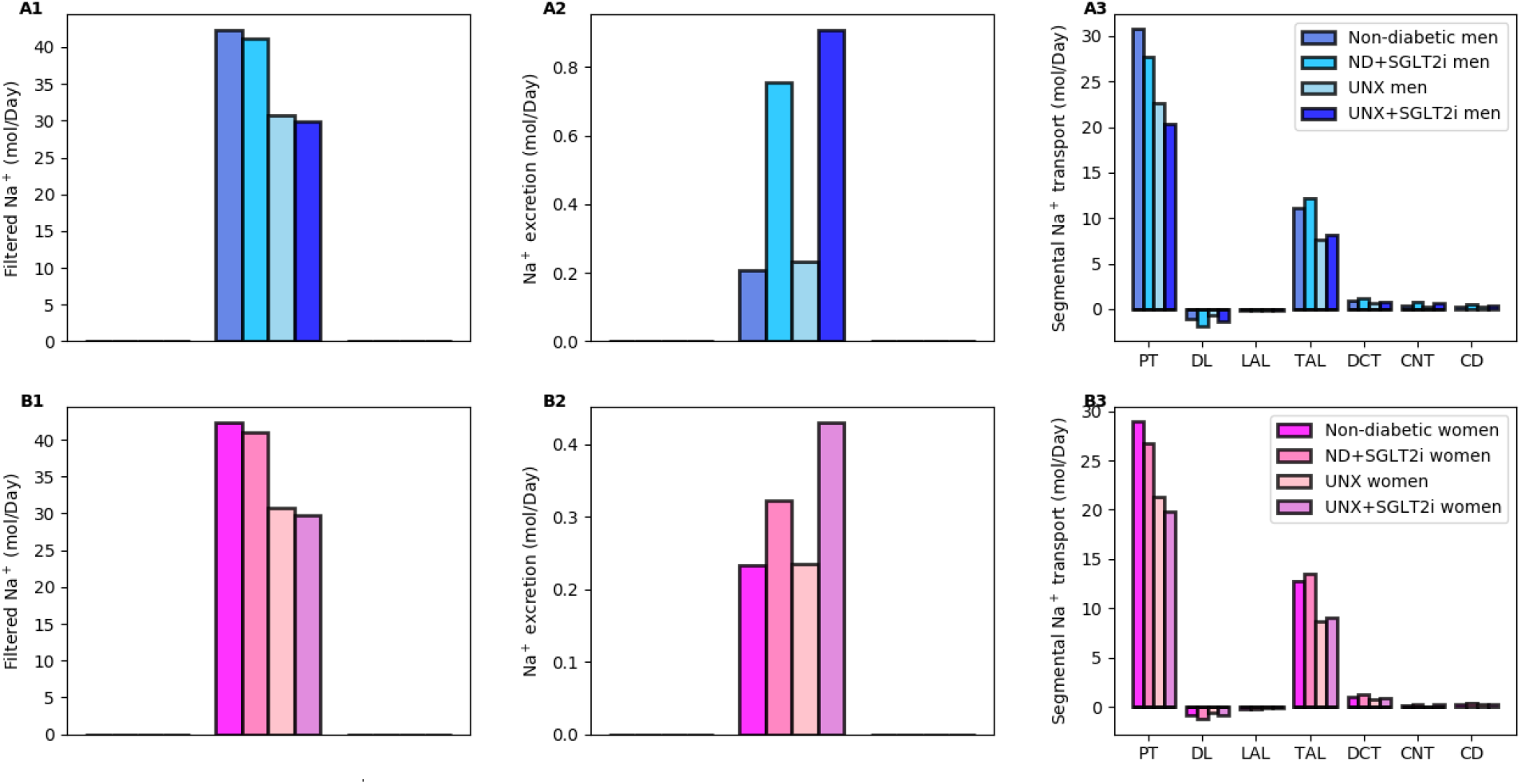
Predicted Na^+^ transport in non-diabetic sham and UNX kidney, in men (top row) and women (bottom row). Results were obtained with and without SGLT2 inhibition. PT, proximal tubule; DL, descending limb; LAL, long ascending limb; TAL, thick ascending limb; DCT, distal convoluted tubule; CNT, connecting tubule; CD, collecting duct.

**Figure 2.**
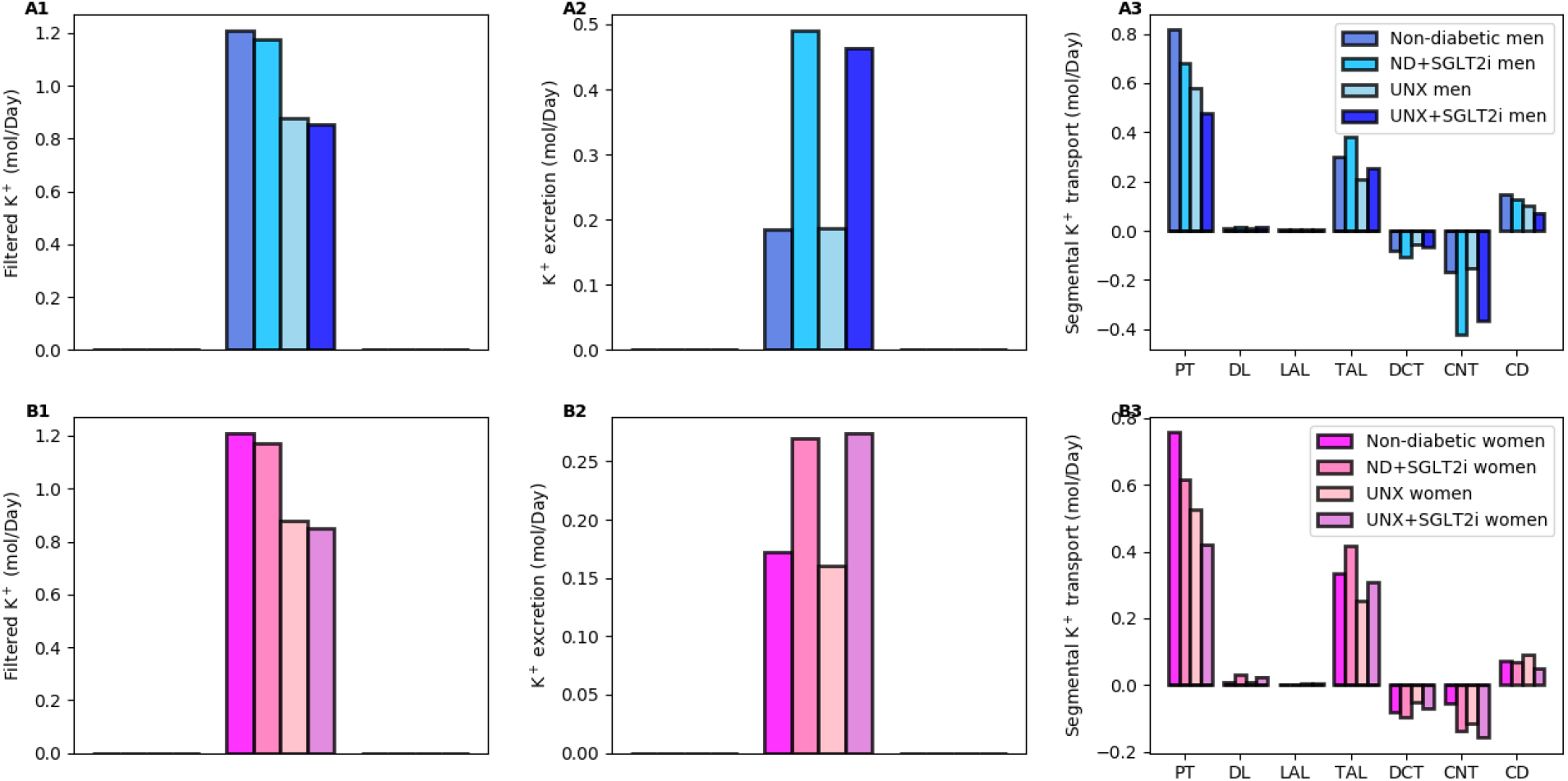
Predicted K^+^ transport in non-diabetic sham and UNX kidney, in men (top row) and women (bottom row). Results were obtained with and without SGLT2 inhibition. Notations are analogous to Fig. 1.

**Figure 3.**
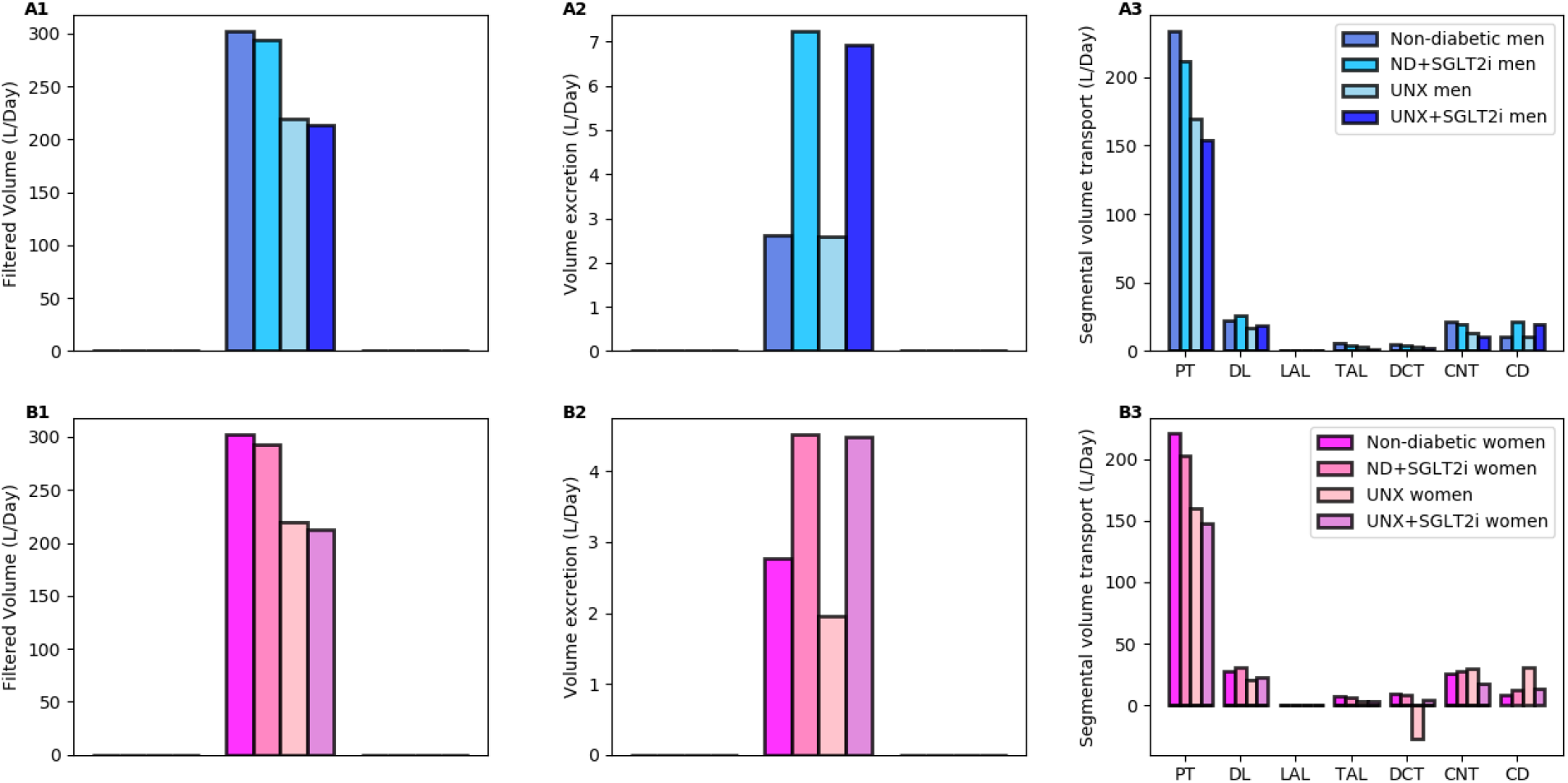
Predicted volume transport in non-diabetic sham and UNX kidney, in men (top row) and women (bottom row). Results were obtained with and without SGLT2 inhibition. Notations are analogous to Fig. 1.

How does nephron loss affect the effects of SGLT2 inhibition? Total filtered glucose load in UNX is 29 and 70% less than sham. However, nephron loss induces hyperfiltration in remaining nephrons; thus single-nephron-filtered glucose load is higher in UNX than sham. Overall, the effects of SGLT2 inhibition in a nondiabetic UNX kidney are qualitatively similar to those for the sham kidney. UNX lowers the glucosuric, diuretic, natriuretic and kaliuretic effect of SGLT2 inhibition (Figs. 1-3).

### Kidney function in diabetic women and men under SGLT2 inhibition

SGLT2 inhibition attenuates diabetes-induced glomerular hyperfiltration and returns GFR to baseline, lowering the filtered glucose load from 1.52 to 1.3 mol·day-1 in moderate diabetes. The kidney’s response in glucose transport is similar in women and men: proximal convoluted tubule glucose reabsorption, mediated by the 10% remaining SGLT2, reduces by ∼80%, from 1.5 to 0.3 mol glucose·day-1. The SGLT1 along the S3 segment reabsorbs a fraction of the remaining glucose, at a rate similar to the proximal convoluted tubule, thereby limiting the risk of hypoglycemia. Glucose excretion in moderately diabetic women and men is predicted to be similar, corresponding to almost 50% of filtered glucose (Figs. 4-6).

**Figure 4.**
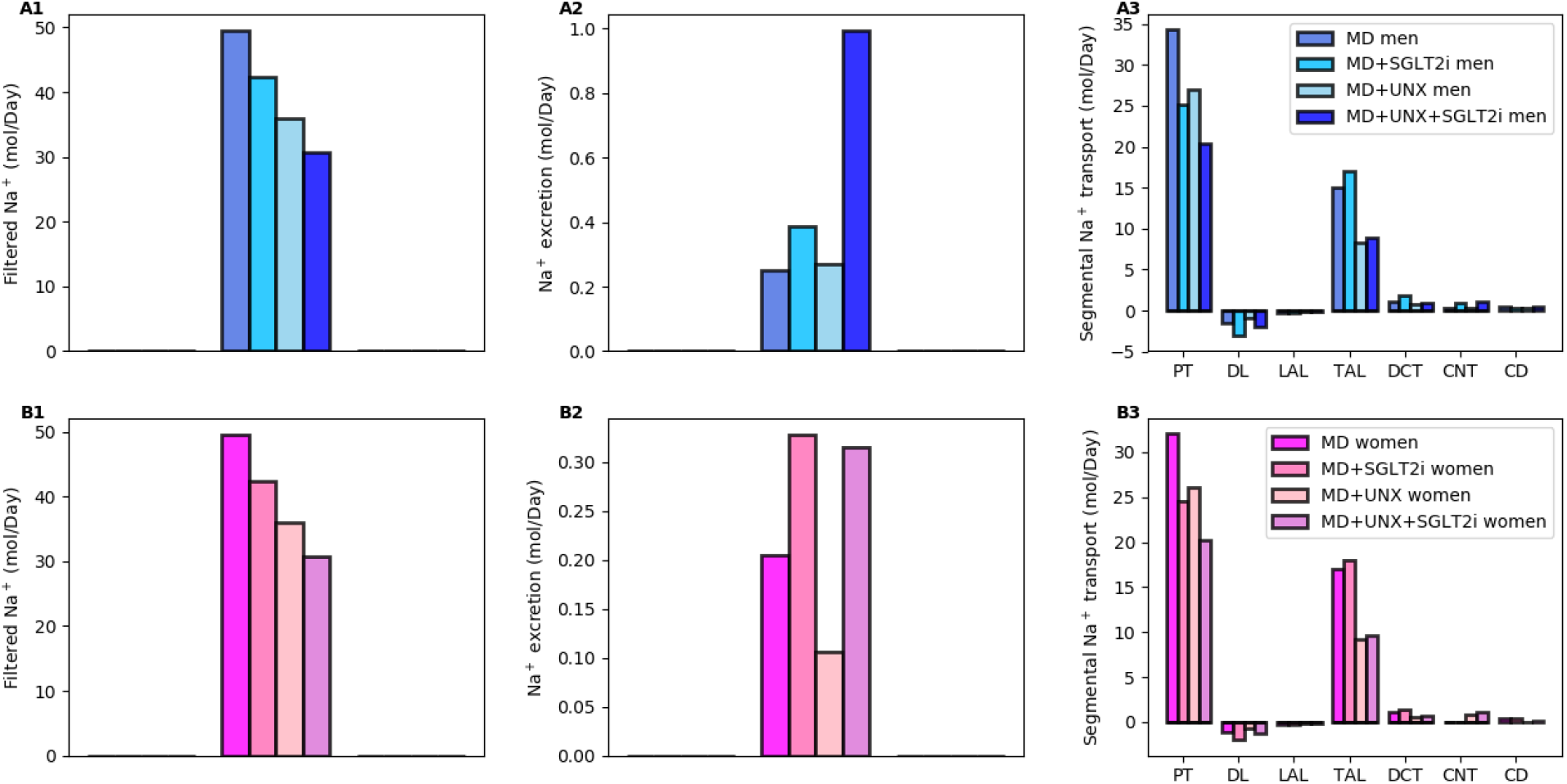
Predicted Na^+^ transport in moderately diabetic sham and UNX kidney, in men (top row) and women (bottom row). Results were obtained with and without SGLT2 inhibition. Notations are analogous to Fig. 1.

**Figure 5.**
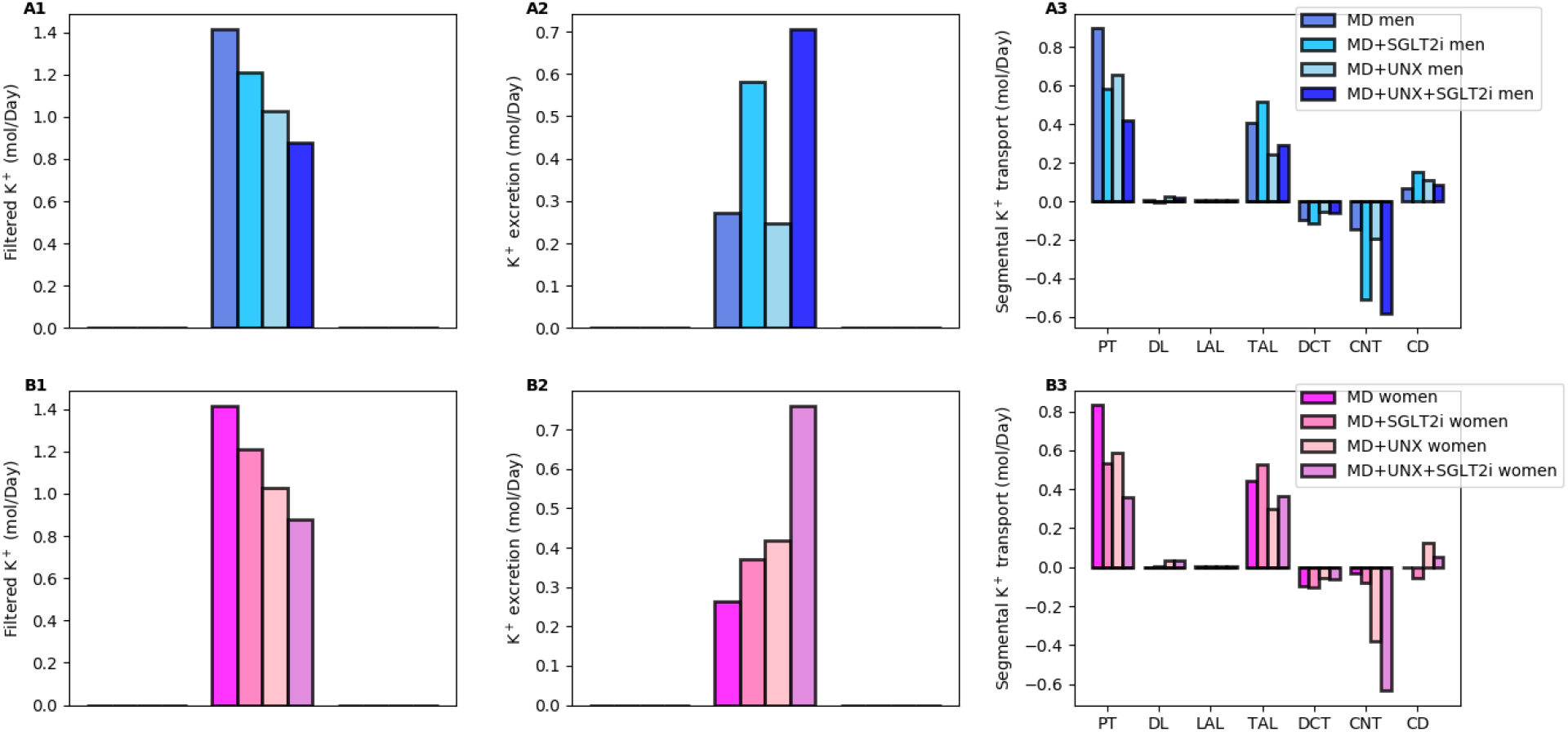
Predicted K^+^ transport in moderately diabetic sham and UNX kidney, in men (top row) and women (bottom row). Results were obtained with and without SGLT2 inhibition. Notations are analogous to Fig. 1.

**Figure 6.**
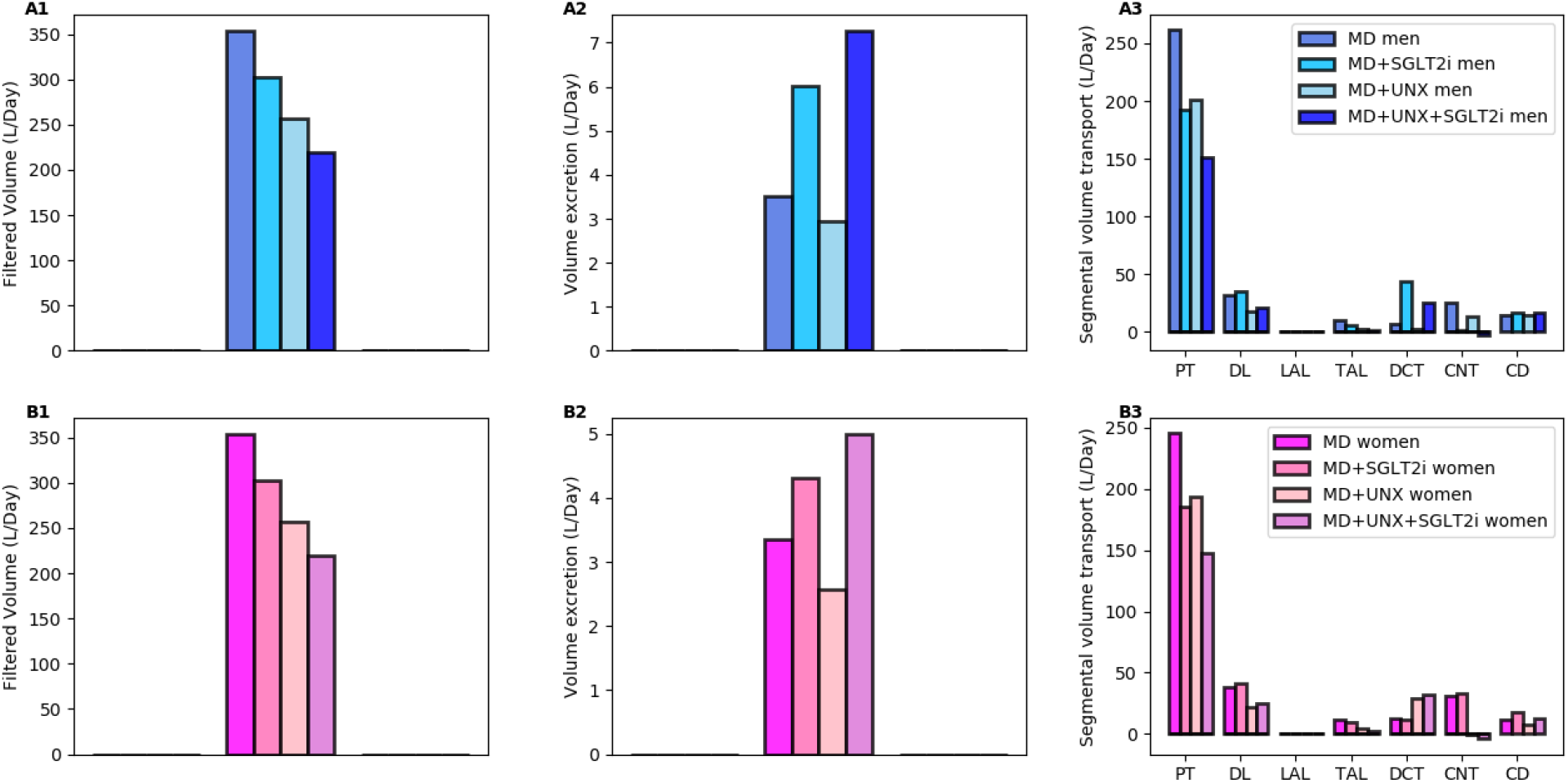
Predicted volume transport in moderately diabetic sham and UNX kidney, in men (top row) and women (bottom row). Results were obtained with and without SGLT2 inhibition. Notations are analogous to Fig. 1.

While the primary target of SGLT2 inhibitors is glucose, they also have a significant impact on renal Na^+^ transport. By normalizing GFR, SGLT2 inhibition significantly lowers filtered Na^+^ load and Na^+^ transport. Our simulations suggest that SGLT2 inhibition induces osmotic diuresis in the proximal tubules, as in the non-diabetic case, thereby reducing paracellular transport. In fact, the model predicts that osmotic diuresis reverses the direction of paracellular Na^+^ transport in S3: the luminal-to-interstitial Na^+^ concentration gradient favors Na^+^ secretion into the lumen via the tight junctions. Consequently, in the moderate diabetes case, Na^+^ excretion increases substantially. Natriuresis is less severe in women because the thick ascending limbs in their kidneys are more capable of compensating for the reduced Na^+^ transport by the proximal convoluted tubules.

The effects of SGLT2 inhibition in the diabetic UNX kidneys are qualitatively similar to sham. The model predicts that with acute SGLT2 inhibition, a reduction in nephron number lowers the glucosuric, diuretic, natriuretic, and kaliuretic effects (Figs. 4-6).

## Discussion

At normal plasma glucose levels, the kidneys reabsorb essentially all the filtered glucose, in a transport process that is mediated the proximal tubules via the SGLT2 and SGLT1, resulting in little glucose excreted in urine. In diabetes, however, either insulin production or sensitivity is impaired, resulting in a rise in blood glucose level. Chronic exposure to elevated blood glucose levels contribute to changes in kidney structures, including tubular hypertrophy and glomerular hyperfiltration observed in diabetes. The activities of SGLT2/SGLT1 are likely modified in patients with diabetes; however, those changes have not been characterized. Nonetheless, when sufficiently high plasma glucose is coupled with glomerular hyperfiltration, the kidney’s glucose transport capacity may be overwhelmed, resulting in the appearance of glucose in urine, which is traditionally considered a hallmark of diabetes.

The goal of this study was to assess the theoretical impact of diabetes and SGLT2 inhibition on solute transport and urinary excretion in kidneys of men and women, with intact and reduced nephron number. Key predictions are:

- In a nondiabetic kidney, SGLT2 inhibition induces glucosuria, diuresis, natriuresis, and kaliuresis, due in large part to reduced reabsorption by the proximal convoluted tubule. When the nephron number is reduced, these effects are lessened, because of the lower filtration.
- In an intact diabetic kidney, acute SGLT2 inhibition induces diuresis, natriuresis, and kaliuresis. UNX attenuates the acute response to SGLT2 inhibition.
- Given the relations between renal fluid and Na^+^ excretion with blood pressure and heart failure, our findings may explain why the blood pressure lowering and heart failure protective effect of chronic SGLT2 inhibition is preserved in DKD patients with reduced GFR.

The nephron models developed in this study can be used as key components in integrative computational models of whole-kidney function^26-39^, in which interstitial fluid composition would be predicted instead of assumed known *a priori*. The resulting models can simulate whole-kidney function in men and women, and with different health status --- a step towards personalized medicine.

